# HippoGenes: A robust workflow for subregional transcriptomic profiling of the human hippocampus

**DOI:** 10.64898/2026.08.04.742581

**Authors:** Alexander Ngo, Sara Larivière, Jessica Royer, Meaghan Smith, Raúl Rodríguez-Cruces, Youngeun Hwang, Lang Liu, Ziv Gan-Or, Alan C. Evans, Andrea Bernasconi, Neda Bernasconi, Sam Audrain, Alexander J. Barnett, Jacob W. Vogel, R. Nathan Spreng, Robert Leech, Sofie L. Valk, Matthias Kirschner, Jordan DeKraker, Boris C. Bernhardt

**Author notes:** Joint senior authors (Jordan DeKraker, Boris C. Bernhardt).

## Abstract

The human hippocampus is a unique cortical structure central to brain function, plasticity, and disease. Unravelling its complex organization requires the integration of multiscale data, linking molecular features to mesoscale anatomy and macroscale functional patterns. Gene expression is a fundamental microscale phenotype, and its profiling can provide a reference description of how molecular features are distributed across the brain. Capitalizing on recent imaging-transcriptomic analyses, we introduce HippoGenes, a repository of fine-grained gene expression patterns across human hippocampal subregions. We leveraged spatial statistical models and hippocampal surface mapping to reconstruct dense transcriptomic maps from sparse *post-mortem* tissue samples of the Allen Human Brain Atlas, generating continuous expression estimates for thousands of genes aligned to a common surface-based coordinate system. We illustrate the utility of HippoGenes to (*i*) map medial-lateral and anterior-posterior transcriptomic gradients that align with subfield and tripartite subdivisions of the hippocampal formation, (*ii*) examine associations between gene expression and canonical microstructural and functional features of the hippocampus, and (*iii*) perform a molecular decoding of subregional alterations in neurological patients with hippocampal pathology. HippoGenes provides a framework for exploring the molecular organization of the hippocampus, opening avenues for multiscale integration in health and disease, and is openly available on https://hippogenes.readthedocs.io/.

## Introduction

The human hippocampus is a complex structure that underpins many aspects of cognition and behaviour. Although traditionally treated as a unitary anatomical entity, converging evidence has established that hippocampal organization is structured along two principal axes. On one hand, medial-to-lateral (transverse) organization reflects cytoarchitectonic and laminar differentiation into canonical subfields, including Cornu Ammonis (CA) 1-4, dentate gyrus, and subiculum, and supports distinct computational roles within hippocampal circuits (Andersen et al., 2006; de Flores et al., 2020; DeKraker et al., 2020, 2026; Duvernoy, 1988; Kulaga-Yoskovitz et al., 2015; Olsen et al., 2019; Paquola et al., 2020; Yushkevich et al., 2015). On the other hand, anterior-to-posterior (longitudinal) organization, spanning the head, body, and tail of the hippocampus, depicts variation in connectivity and functional specialization. Anterior regions are strongly implicated in affective processing, associative and semantic memory, and integration with lateral temporal and limbic regions, while posterior regions are closely linked to spatial cognition, contextual representation, and sensory detail (Borne et al., 2023; Genon et al., 2021; Poppenk et al., 2013; Strange et al., 2014). The interaction between these axes reflects a coordinated structure-function relationship, whereby continuous longitudinal gradients intersect with discrete subfield-specific circuits across spatial scales (Daugherty et al., 2026; DeKraker et al., 2025; Duvernoy, 1988; Genon et al., 2021; Knierim & Neunuebel, 2016; Strange et al., 2014; Vos de Wael et al., 2018). Achieving a comprehensive understanding of hippocampal organization therefore requires a multiscale, integrative framework: from macroscale hierarchies of connectivity and function, through mesoscale structure, down to microscale molecular features reflected in transcriptomic patterns.

Open release of *postmortem* human transcriptomic datasets, such as the Allen Human Brain Atlas (AHBA) with over 20,000 genes across spatially distinct tissue samples, have made it possible to explore gene expression patterns in the brain (Arnatkevic iūtė et al., 2019; Hawrylycz et al., 2012). This provides a molecular signature shaping cellular composition, circuit organization, and regional specialization (Hawrylycz et al., 2012; Lein et al., 2007; Tasic et al., 2018). Studies leveraging AHBA have characterized large-scale transcriptomic variation across the neocortex, revealing coherent molecular hierarchies aligned with sensory-association axes, developmental trajectories and macroscale connectivity patterns (Burt et al., 2018a; Markello et al., 2022; Vogel et al., 2024; Wagstyl et al., 2024). Despite these advances, the hippocampus is comparatively underrepresented, often summarized as a single region-of-interest where data are averaged across the entire structure or subfields. Consequently, the molecular organization of the hippocampus remains unexplored at spatial scales that parallel its fine-grained subregional architecture.

Resolving fine-grained gene expression patterns within the hippocampus is challenging due to its folded topography and the inherently sparse, uneven sampling of *post-mortem* tissue. However, brain transcriptional profiles are not spatially random, but rather exist in a spatially embedded system, such that nearby regions exhibit greater similarity than more distant ones (Cembrowski & Spruston, 2019; Krienen et al., 2016; Richiardi et al., 2015; Strange et al., 2014; Vértes et al., 2016). This spatial autocorrelation arises, in part, from neurobiological constraints including shared developmental origins, continuity of cytoarchitecture, and cell-type composition, and locally structured connectivity, which together give rise to smooth transcriptional gradients across anatomical space (Burt et al., 2018b; Hawrylycz et al., 2012; Vogel et al., 2024). These patterns provide a theoretical foundation for modelling approaches that leverage distance-dependent relationships to infer expression at unsampled locations. One such approach is kriging, an interpolation technique that predicts values at unsampled locations by weighting nearby observations according to their spatial similarity and distance relationships. By modelling both broad spatial trends and local variation, kriging enables the reconstruction of continuous, vertex-wise maps from sparse transcriptomic samples (Ecker et al., 2025; Gryglewski et al., 2018; Unterholzner et al., 2020; Wackernagel, 1995). Combined with surface-based representations of the hippocampus, these approaches offer a principled means to infer microscale molecular information at degrees of granularity otherwise inaccessible from sparse transcriptomic sampling, while simultaneously linking it to mesoscale anatomy and macroscale organization (Bernhardt et al., 2016; DeKraker et al., 2022, 2025; Royer et al., 2026).

Here, we introduce HippoGenes, an open-access, online data repository for hippocampal gene expression (https://hippogenes.readthedocs.io/). Using sparse *post-mortem* tissue samples from AHBA and donor-specific surface representations of the hippocampus derived from the corresponding MRI, we employed shape-intrinsic alignment to map sample locations to a standard coordinate system. Spatial statistical models (kriging) were applied to interpolate transcriptional data reconstruct a vertex-wise atlas of hippocampal expression for 13,651 genes. To demonstrate the utility of reconstructed, vertex-wise atlas of gene expression, we (*i*) studied transcriptional organization of the hippocampus and its associations with medial-lateral and anterior-posterior subdivisions, (*ii*) contextualize these gene expression patterns with microstructural and functional features, and (*iii*) performed hypothesis-guided gene interrogation in disease, particularly in patients with epilepsy. Together, this framework provides a resource for exploring the molecular organization of the hippocampus, opening avenues for multiscale integration and associations with health and disease.

## Results

### Transcriptional mapping of the human hippocampus

To establish spatially continuous molecular representations of the hippocampus, we reconstructed a vertex-wise atlas of hippocampal expression for 13,561 genes using a sparse-to-dense interpolation of anatomically localized *post-mortem* samples (**Figure 1**).

**Figure 1.**
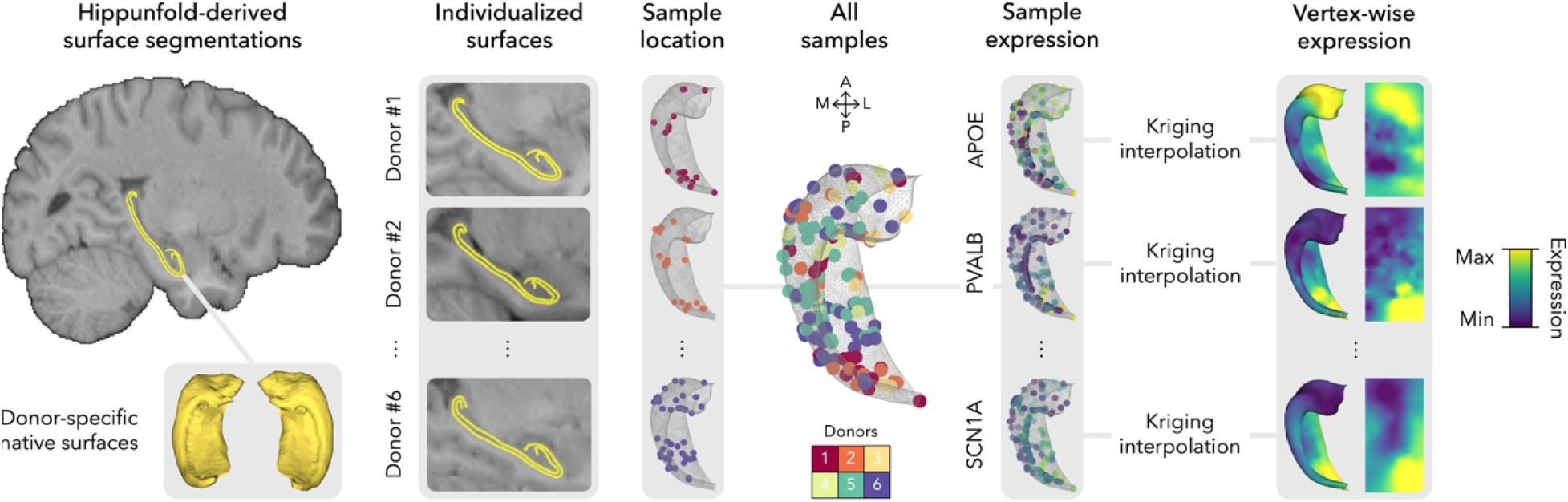
**(A)** Schematic depiction of sparse-to-dense mapping of hippocampal samples using spatial autocorrelation preserving interpolation.

Cross-validation demonstrated stable and reproducible spatial gene expression patterns. In leave-one-sample-out analyses, gene-wise expression profiles estimated after withholding individual samples remained highly similar (mean ± standard deviation [SD] correlation = 0.967 ± 0.009; range = 0.907 – 0.978; **Supplementary figure 1**), indicating that no single sample disproportionately influenced the resulting atlas. Consistent with this observation, leave-*k*-out analyses (*k* = 10-90% of samples withheld, in 10% increments) recovered expression patterns that remained strongly correlated with the full-data atlas. Although correlation strength progressively decreased as sampling density reduced, reconstruction performance did not linearly scale with the number of held-out samples, indicating reliable approximation spatial gene profiles (**Supplementary table 1**; **Supplementary figure 2**).

### Transcriptomic pathways of the medial-lateral and anterior-posterior hippocampal axes

Surface-based gene atlas enables data-driven gene interrogation of any hippocampal feature. We employed partial least squares (PLS) to identify sets of genes that are systematically expressed along the principal anatomical axes of the hippocampus, revealing significant latent variables whose spatial expression patterns align with the geodesic medial-lateral and anterior-posterior coordinates of the hippocampus.

Along the medial-lateral axis, the first latent variable significantly explained 83.7% of the covariance between geodesic transversal coordinates and gene expression (*p*_perm_ < 0.05). Gene set enrichment analysis of this gene set identified processes involved in ribosomal assembly, cell components, RNA binding, synaptic function, and neuron generation (all *p*_FDR_ < 0.05), which may be related to cell division, differentiation, and proliferation (**Figure 2A**).

**Figure 2.**
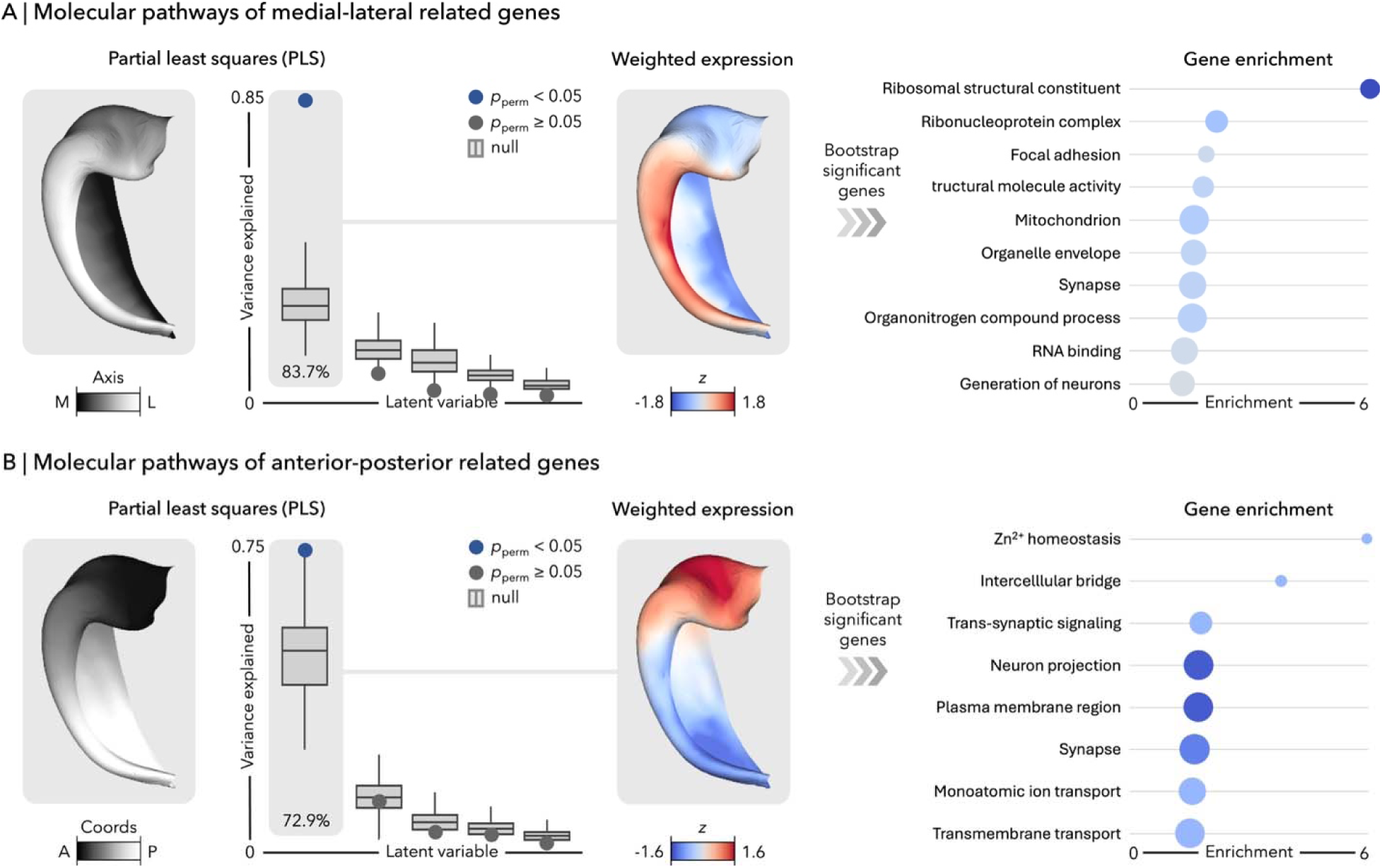
**(A)** Partial lease squares (PLS) analysis linking the geodesic transverse (medial-lateral) axis of the hippocampus to spatially dense gene expression patterns. *Left*: Variance explained by latent variables compared to permutation-based null models. In the boxplots, the ends of box represent the first (25%) and third (75%) quartiles, the centre line (median) represents the second quartile of the null distribution (1000 permutations), the whiskers represent the non-outlier endpoints of the distribution. Points represent empirical observations (with significance defined as *p*_perm_ < 0.05). *Middle*: Spatial distribution of weighted gene expression of first latent variable (LV1). *Right*: Gene ontology enrichment of top 5% of bootstrap-significant genes. Each ontology term represents a biological pathway or cellular process defined by a set of genes. Circle size represents the number of genes that overlap in the given term and colors indicate FDR-corrected p-value (-log_10_(*p_F_*_DR_)). *Abbreviations*: L: lateral; M: medial; *p*_perm_: *p*-value based on permutation tests controlling for spatial autocorrelation. **(B)** Repeated analyses for geodesic longitudinal (anterior-posterior) axis of the hippocampus. *Abbreviations*: A: anterior; M: posterior.

In a separate PLS, covariation between anterior-posterior axis and gene expression was explainable by 72.9% in the first latent variable (*p*_perm_ < 0.05). These genes reflected processes associated with synaptic activity and communication as well as neuronal projection (all *p*_FDR_ < 0.05; **Figure 2B**).

### Gene expression organization and its relation to multiscale features of the hippocampus

Studying the intrinsic organization of transcriptomic profiles within the hippocampus, non-linear dimensionality reduction was used to summarize the information contained across the co-expression of hippocampal regions. This revealed two principal eigenvectors describing the spatial topography of variation. The primary axis (G1), which explained 27.5% of the variance, predominantly followed the main medial-lateral differentiation of the hippocampus (*r* = 0.84, *p*_perm_ < 0.05), and the secondary axis (G2), which accounted for 15.7%, mainly captured an anterior-posterior organizational pattern (*r* = −0.51, *p*_perm_ < 0.05; **Figure 3A**).

**Figure 3.**
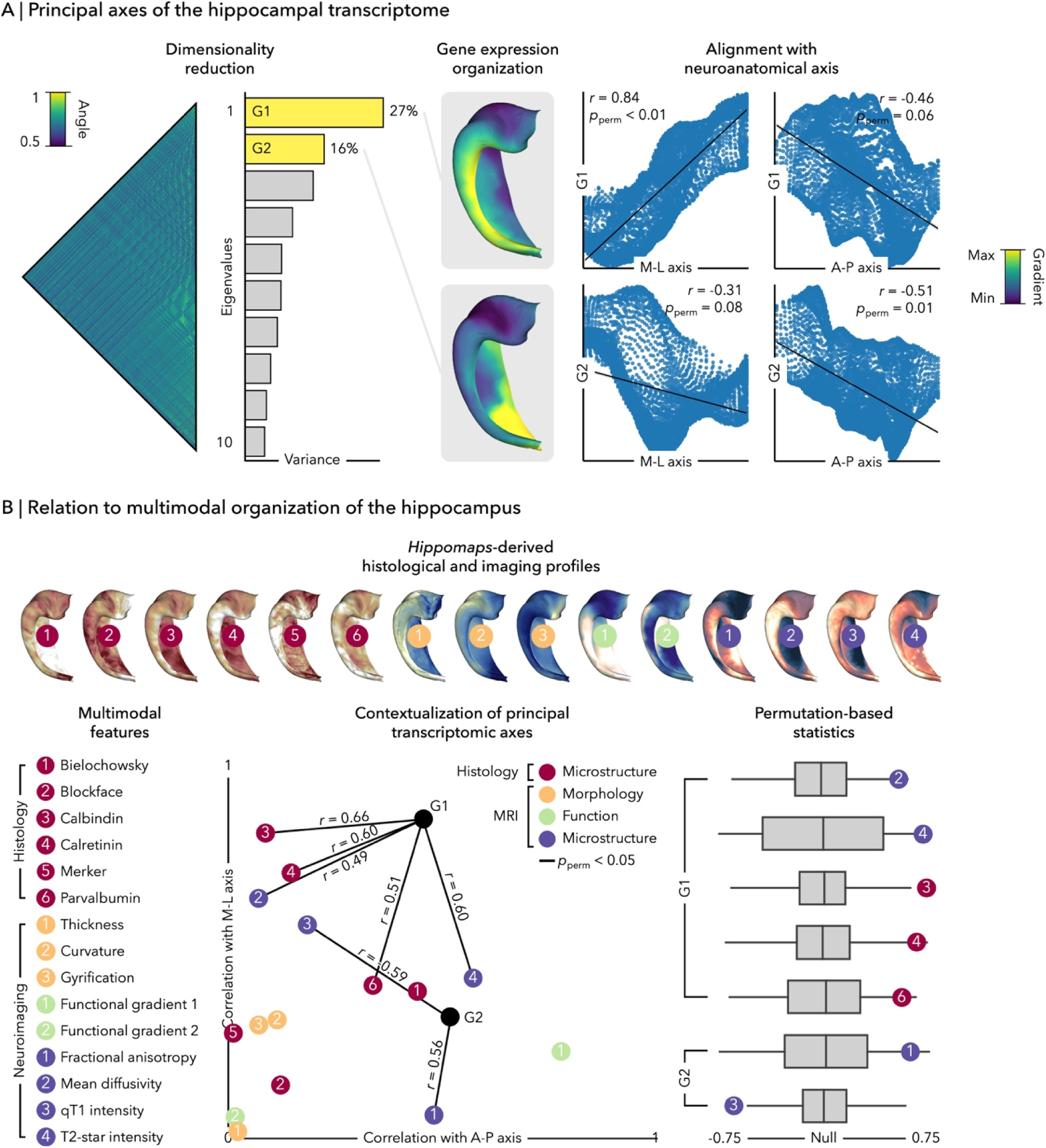
**(A)** Organization of gene expression within the hippocampus. *Left*: Latent representations of gene coexpression profiles across different regions. *Middle*: Spatial distribution of the first two eigenvectors from non-linear dimensionality decomposition (G1 and G2). Colours represent the coordinate of a vertex along a transcriptomic axis identified by dimensionality reduction, where similar values indicate similar transcriptomic profiles. *Right*: Correlation between eigenvectors and anatomical medial-lateral and anterior-posterior axes. *Abbreviations*: A-P = anterior-posterior; G1 = primary axis of transcriptomic organization; G2 = secondary axis of transcriptomic organization; M-L = medial-lateral. *p*_perm_: *p*-value based on permutation tests controlling for spatial autocorrelation. **(B)** Alignment with hippocampal multiscale and multimodal hierarchies. *Top*: *HippoMaps*-derived profiles (Red: Bieloschowsky [1], blockface [2], calbindin [3], calretinin [4], Merker [5], and parvabumin [6]; yellow: curvature [1], gyrification [2], and thickness [3]; green: resting-state functional gradient one [1] and two [2]; purple: fractional anisotropy [1], mean diffusivity [2], qT1 intensity [3], and T2-star intensity [4]). *Bottom left*: List of features. *Bottom middle*: Absolute correlation between each feature map and the geodesic anterior-posterior and medial-lateral coordinates of the hippocampus. Solids lines represent significant spatial correlation between feature map and transcriptomic axes (G1 and G2 from Figure 3A). *Bottom right*: Significant spatial correlations compared against permutation-based null models. In the boxplots, the ends of box represent the first (25%) and third (75%) quartiles, the centre line (median) represents the second quartile of the null distribution (1000 permutations), the whiskers represent the non-outlier endpoints of the distribution. Points represent empirical observations.

To situate the principal axes of gene expression within the broader organizational architecture of the hippocampus, cross-modality correlational analyses quantified the association between the main spatial patterning of gene expression and established functional, microstructural, and cellular properties derived from *post-mortem* and *ex-vivo* histology as well as *in-vivo* structural and functional MRI (**Figure 3B**; DeKraker et al., 2025).

Systematic spatial correlation analysis demonstrated that the primary gene axis (G1; see *Figure 4A*) showed strongest correspondence with microstructural properties (mean diffusivity and T2-star intensity) as well as inhibitory interneuron composition (calbindin, calretinin, and parvalbumin; all *p*_perm_ < 0.05). The secondary axis (G2; see *Figure 4A*) demonstrated preferential associations with intracortical myeloarchitecture and microstructure (qT1 intensity and fractional anisotropy; all *p*_perm_ < 0.05). These predominant patterns of transcriptomic differentiation capture biologically meaningful organization dimensions of the hippocampus that extend across functional microstructural, and cellular scales (**Figure 3B**).

**Figure 4.**
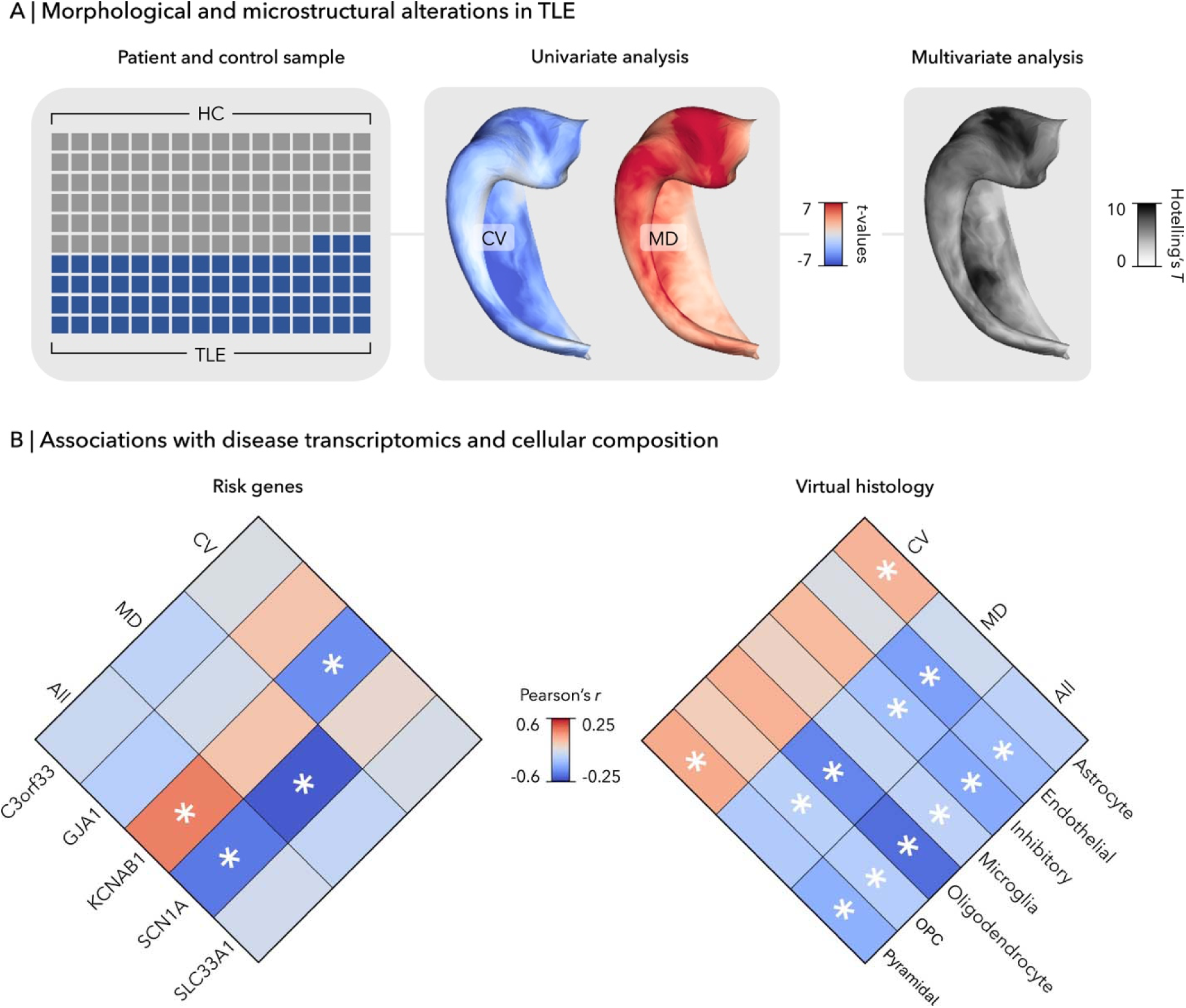
**(A)** Structural changes in the hippocampus of patients with temporal lobe epilepsy (TLE). *Left*: Schematic representation of the sample, with each square corresponding to an individual participant (grey for controls and blue for patients). *Middle*: Vertex-wise univariate case-control comparisons showing differences in columnar volume (CV) and mean diffusivity (MD) between patients and controls. Blue and red colours represent decreases and increases in TLE. *Right*: Multivariate case-control differences obtained by jointly modeling CV and MD. Black colours represent increased alterations in patients. **(B)** Spatial correlation between TLE-(*left*) and cell-specific (*right*) genes and univariate and multivariate changes in hippocampus. Statistical significance (asterisk) was assessed using two-tailed, non-parametric tests controlling for spatial autocorrelation. *Abbreviations*: CV = columnar volume; HC = healthy controls, MD = mean diffusivity; OPC = oligodendrocyte precursor cell; TLE = temporal lobe epilepsy.

### Disease risk associations with regional hippocampal vulnerability

To illustrate the clinical relevance of hippocampal gene expresison, we assessed its relationship with structural vulnerability in temporal lobe epilepsy (TLE), a focal epilepsy classically associated with hippocampal pathology. We focused on hippocampal volume (CV) and mean diffusivity (MD), as these complementary MRI-derived markers provide comprehensive assessment of disease-related pathology through macroscopic tissue loss and microstructural tissue integrity. In line with prior literature (Bernhardt et al., 2016; Caldairou et al., 2021; Cendes et al., 1993; Ripart et al., 2025), patients with TLE exhibited widespread patterns of ipsilateral structural compromise compared to healthy controls, characterized by regional hippocampal atrophy and increased mean diffusivity (**Figure 4A**).

Among *a priori* genes previously linked to TLE (C3orf33, GJA1, KCNAB1, SCN1A, SLC33A1; Abou-Khalil et al., 2018; Kasperavičiūtė et al., 2013), regional atrophy was strongly associated with KCNAB1 expression (*r* = −0.41, *p*_perm_ < 0.05) and microstructural alterations with SCN1A expression (*r* = −0.56, *p*_perm_ < 0.05), highlighting genes implicated with neuronal excitability and seizure susceptibility (**Figure 4B**).

Beyond individual risk genes, spatial pattern of TLE-related alterations also exhibited distinct cellular profiles. Atrophy preferentially localized to regions enriched for pyramidal neuron and astrocytic processes (range median *r* = 0.09 to 0.10, all *p*_perm_ < 0.05). However, changes in microstructure were associated with oligodendrocyte precursor cells (OPC), oligodendrocytes, inhibitory neurons, and endothelial cells (range median *r* = −0.06 to −0.18, all *p*_perm_ < 0.05; **Figure 4B**), which are closely linked to tissue microenvironment, myelination and neurovascular organization.

## Discussion

Despite substantial progress in aggregating and disseminating multiscale features of the human hippocampus derived from neuroimaging and histology (DeKraker et al., 2025), we currently lack a unified anatomy-driven framework for mapping its molecular architecture and embedding it within subregional hippocampal organization. Leveraging advances in hippocampal computational anatomy, imaging-transcriptomics and spatial statistical modeling, we establish a comprehensive, surface-based atlas of the hippocampal transcriptome, HippoGenes. This is designed to (*i*) characterize fine-grained gene expression patterns, (*ii*) integrate molecular architecture across multimodal and multiscale domains, and (*iii*) enable data-driven discovery and hypothesis-guided gene interrogation in both health and disease. By bridging precision gene mapping with systems neuroscience, HippoGenes offers a roadmap for decoding the hippocampus at an unprecedented scale, enabling mechanistic understanding from molecular specialization to network-level function.

Advances in high-throughput transcriptomic profiling and spatial analytical methods have enabled the construction of anatomically comprehensive gene expression atlases of the human brain, fueling the rapid growth of imaging-transcriptomic applications (Hawrylycz et al., 2012). These approaches have uncovered transcriptional correlates of a diverse range of structural and functional properties of the brain, and have illuminated the molecular substrates of brain alterations observed across neurological and psychiatric conditions (Arnatkeviciute et al., 2022; Markello et al., 2021). While these analyses have predominantly been performed in the neocortex at different spatial resolutions ranging from coarse parcel to vertex-wise maps (Ecker et al., 2025; Gryglewski et al., 2018; Larivière et al., 2021; Markello et al., 2021; Unterholzner et al., 2020; Wagstyl et al., 2024), transcriptomic investigations of the human hippocampus have often relied on a region-of-interest approach, averaging expression values across the entire structure or subfields (Kember et al., 2025; Larivière et al., 2021, 2022). Such strategies may obscure fine-grained spatial variation, an organization feature consistently observed in hippocampal histology and structural and functional imaging (DeKraker et al., 2025; Duvernoy, 1988; Olsen et al., 2019). Recent methodological developments have overcome this limitation, notably surface-based reconstruction techniques that preserve hippocampal geometry and subfield topology within a standardized unfolded coordinate system (Bernhardt et al., 2016; Caldairou et al., 2016; DeKraker et al., 2021, 2022, 2023). Within this framework, sparse *post-mortem* tissue samples can be projected onto a common surface representation and subsequently interpolated. In HippoGenes, the application of ordinary kriging leverages the empirically observed spatial autocorrelation of gene expression to estimate values at unsampled regions as the weighted combination of neighbouring observations (Wackernagel, 1995). By explicitly modeling the spatial covariance structure, this captures both large-scale and local transcriptional variation while minimizing estimation bias. Similar to prior methodological research in the cortex (Ecker et al., 2025; Gryglewski et al., 2018; Unterholzner et al., 2020), this sparse-to-dense reconstruction transforms discretely sampled microarray measurements into continuous, vertex-wise expression maps that are directly comparable to surface-based histology and imaging analyses. Cross-validation analyses confirmed the consistency and reproducibility of these reconstructed profiles, establishing a robust molecular atlas. By providing continuous surface-based maps of gene expression, the atlas enables direct integration of transcriptomic, histological, and neuroimaging data within a common spatial space. This approach facilitates the discovery of genes and molecular pathways associated with spatial variation in hippocampal structure and function.

The hippocampus is characterized by a highly structured architecture spanning multiple spatial axes, most prominently medial-lateral and anterior-posterior axes (Genon et al., 2021). These dimensions reflect coordinated differences in cytoarchitecture, connectivity, and functional specialization, forming a graded organizational framework rather than sharply defined boundaries between regions (Angeli et al., 2025; DeKraker et al., 2022; Duvernoy, 1988; Genon et al., 2021; Kharabian Masouleh et al., 2020; Paquola et al., 2020; To et al., 2025; Vos de Wael et al., 2018). Consistent with this principle, large-scale molecular variation along the hippocampus also follows continuous spatial gradients, suggesting that molecular architecture is aligned with these fundamental organizational axes (Vogel et al., 2020). The dominant transcriptomic pattern primarily reflects medial-to-lateral differentiation consistent with differences in subfield microstructure and cellular composition. Given the well-documented variations in cytoarchitecture and connectivity patterns (Amunts et al., 2005; Ding & Van Hoesen, 2015a, 2015b; Palomero-Gallagher et al., 2020), different subfields also exhibit distinct genetic and transcriptomic profiles that support their molecular specialization (Ianov et al., 2017; Lein et al., 2004; Thompson et al., 2008; van der Meer et al., 2020; X. Zhao et al., 2001). Medial-to-lateral expression of genes associated with synaptic function as well as pathways regulating cell division, differentiation, and proliferation may therefore contribute to the establishment and maintenance of cytoarchitectural boundaries, supporting different functional roles of hippocampal subfields. In parallel, complimentary transcriptomic axis (G2) predominantly captures variation along the anterior-posterior axis of the hippocampus, a principal organizational dimension consistently linked to brain function and large-scale network integration (Poppenk et al., 2013; Strange et al., 2014). The underlying subfield arrangement and intrinsic circuitry of the hippocampus are largely preserved along this axis, including across species, supporting the presence of a relatively stereotyped computational microcircuit at each longitudinal position (DeKraker et al., 2024). Given this conservation of local circuitry, the anterior–posterior axis is nevertheless associated with pronounced differences in connectivity and functional specialization. This dissociation suggests that longitudinal variation may not reflect distinct computations, but rather differences in the inputs and outputs that shape the content and context of hippocampal processing (Borne et al., 2023; Poppenk et al., 2013; Przeździk et al., 2019; Vos de Wael et al., 2018). Consistent with this framework, genes that are differentially expressed along this axis are enriched for molecular pathways related to synaptic activity, intercellular communication, and neuronal projection, supporting processes underlying information transfer and long-range connectivity. This representation aligns with well-established brain systems, whereby anterior hippocampal regions are embedded, both transcriptionally and functionally, within networks including medial prefrontal and anterior temporal areas that support affective, social, and semantic processing, while posterior regions are closely aligned with occipital, parietal, and sensorimotor systems involved in contextual, spatial and perceptual functions (Borne et al., 2023; Genon et al., 2021; Poppenk et al., 2013; Vogel et al., 2020; Xie et al., 2024). Large-scale molecular organization may be anchored to the canonical medial-lateral and anterior-posterior hippocampal axes, linking transcriptional variation to different multiscale organizational properties in both health and disease.

While many neurological and psychiatric disorders targeting the hippocampus are often characterized as affecting the structure globally, their impact is rarely uniform. Pathological alterations tend to exhibit distinct patterns across different hippocampal regions. In TLE, atrophy and microstructural disruption are most pronounced in certain subfields and in anterior portion of the hippocampus (Bernasconi et al., 2003; Bernhardt et al., 2016; Blümcke et al., 2013; Caldairou et al., 2021; Goubran et al., 2015; Larivière et al., 2024; Steve et al., 2020). This patterned vulnerability was aligned to the spatial expression of disease-relevant risk genes, most notably KCNAB1 and SCN1A. These genes regulate ion channel function and neuronal excitability through potassium and sodium channel dynamics (Liu & Bean, 2014; Matricardi et al., 2023). Intrinsic regional differences may confer susceptibility to seizure-related structure damage, through hippocampal hyperexcitability, impaired inhibitory regulation, neuronal injury, and gliosis associated with epileptic activity (Chen et al., 2025; Kasperavičiūtė et al., 2013; Oyrer et al., 2018; Wang et al., 1994; Wenzel et al., 2007). Beyond individual epilepsy-related genes, alterations were also linked to the expression of genes enriched in specific cellular populations. Differences in cellular composition and microenvironmental support may also shape disease expression. Atrophy in regions enriched for pyramidal neuron markers is consistent with hippocampal sclerosis, while diffusion changes associated with variation in glial and vascular signatures may reflect contributions from related neuroinflammatory and metabolic processes (Blümcke et al., 2013; Goyal et al., 2025; Stefanits et al., 2012). Despite a lack of literature surrounding the direct associations between regional transcriptomic profiles and hippocampal vulnerability, linking macroscale imaging phenotypes with underlying molecular and cellular organization provides an important framework for future investigations identifying candidate genes and biological pathways that may support, mediate, or protect against hippocampal alterations in TLE. While it was an exemplary disease application of hippocampal transcriptomics, the utility of this framework can extend beyond epilepsy. Many neurological and psychiatric disorders exhibit non-uniform patterns of hippocampal involvement that preferentially affect specific subfields and axes of organization (de Flores et al., 2015; Jubeir & Jacob, 2026; Small et al., 2011; Sun et al., 2023). Spatially resolved maps of gene expression and cellular composition can be used to identify genes, pathways, and cell populations that may coincide with regions of heightened alterations, generating mechanistic hypotheses and advancing our understanding of different pathologies.

The hippocampal gene expression atlas relies on data from AHBA, which represents the most comprehensive publicly available resource for spatially resolved human brain gene expression, but several limitations should be acknowledged. It is comprised of a relatively small number of donors, with uneven hemispheric sampling, demographic variability, and differences in *post-mortem* intervals, all of which may influence expression values despite extensive normalization procedures (Arnatkeviciute et al., 2022). While we mitigate some sources of anatomical variability using accurate subject-specific hippocampal segmentation and surface-based alignment, cross-donor pooling still precludes the assessment of inter-individual differences, including effects of genotype or environment, and may obscure biologically meaningful variability (Grishkevich & Yanai, 2013; Johansen et al., 2023). This is particularly relevant when attempting to relate genetic findings to transcriptional patterns. In addition, sample-level expression profiles are derived from microarray-based measurements, a cost-effective and widely used approach for transcript profiling. This is restricted to preselected probes and may therefore miss lowly expressed or unannotated transcripts. Interpretation of sample expression values is further constrained as probe intensity provides an indirect estimate of transcript abundance and may be affected by platform-specific noise and signal limitations. More recent approaches such as RNA sequencing (RNA-Seq) provided broader transcriptome coverage and improved sensitivity. With single-cell and single-nucleus RNA-seq that enable more refined characterization of cell-type specific expression profiles, this might mitigate limitations inherent to bulk tissue measurements that reflect heterogeneous mixtures of cell populations (Stegle et al., 2015; S. Zhao et al., 2014). Although there is a lack of available RNA-Seq atlases with dense sampling and full hippocampal coverage required for vertex-wise mapping of gene expression, future replication of our findings in larger and more spatially resolved datasets holds significant promise for generating more accurate and detailed transcriptomic maps. Finally, given the nature of imaging-transcriptomics, analyses are inherently correlational and do not provide causal inference. Finally, as with all imaging-transcriptomic analyses, observed associations between gene expression patterns and imaging-derived measures are correlational and do not establish causal mechanisms. These reflect indirect relationships or shared underlying factors rather than direct effects of transcriptional programs on brain phenotypes. Nonetheless, integrating transcriptomics with neuroimaging still provides a valuable framework for generating biologically informed hypotheses, which can be further evaluated through experimental, longitudinal, genetic, and multimodal approaches (Royer et al., 2026).

Overall, the integration of external datasets to contextualize hippocampal structure and function has rapidly advanced in recent years. The increasing availability of open-access resources has enabled systematic investigation of multiscale hippocampal organization in both health and disease. HippoGenes extends this landscape by introducing a spatially dense, surface-based atlas of hippocampal gene expression, adding a critical molecular dimension to existing frameworks. This resource facilitates broad accessibility and continued development through openly available neuroinformatics tools, comprehensive online documentation, and user-friendly workflows designated to support integration with existing hippocampal imaging pipelines and toolboxes. By enabling fine-grained mapping and multiscale integration of transcriptomic profiles, HippoGenes provides a versatile platform for exploring biological foundations of hippocampal organization, identifying candidate genes, pathways, and cellular pathways, as well as disease-related alterations.

## Material and methods

### Allen Human Brain Atlas

Human imaging and gene expression data from six neurotypical adult whole brains (two African American males, three Caucasian males, and one Hispanic female; age range = 24 – 57 years; mean ± SD age = 42.5 ± 13.4 years; **Table 1**) were obtained from the Allen Institute for Brain Science (Hawrylycz et al., 2012). Briefly, tissue samples were extracted across both hemispheres of two human brain donors, as well as the left hemisphere of four additional donors, totaling 3,702 samples. Each sample underwent microarray analysis and preprocessing to quantify gene expression across 58,692 probes, providing an estimate of relative expression of different transcripts (encoded by different genes) within the tissue sample. More in-depth description of the dataset can be found elsewhere (Hawrylycz et al., 2012).

**Table 1.** Demographic data of donors from Allen Human Brain Atlas.

| ID | Age | Sex | Ethnicity | Samples | Hippocampal samples |
| --- | --- | --- | --- | --- | --- |
| 1 | 24 | M | African American | 438 / 458 | 31 / 30 |
| 2 | 39 | M | African American | 433 / 409 | 20 / 28 |
| 3 | 57 | M | Caucasian | 353 / – | 20 / – |
| 4 | 31 | M | Caucasian | 518 / – | 24 / – |
| 5 | 55 | M | Caucasian | 415 / – | 13 / – |
| 6 | 49 | F | Hispanic | 436 / – | 12 / – |

Age is reported as years. Number of tissue samples and subset of hippocampal samples are indicated separately for left and right hemispheres. Abbreviations: M = male; F = female; L = left; R = Right

### Data processing

*i) Imaging processing*. Individualized hippocampal surfaces were reconstructed from donor-specific structural T1-weighted magnetic resonance imaging (MRI) images using *HippUnfold version 1.3.0* (https://hippunfold.khanlab.ca/; DeKraker et al., 2022). This involved tissue type segmentation using a deep UNet neural network, fitting of inner, outer, and midthickness surfaces to hippocampal grey matter, mapping to a standardized unfolded space, and registration in unfolded space to a standard, histology-derived generated atlas (DeKraker et al., 2022, 2023).
*ii) Microarray expression processing*. Probe-level measures of gene expression for all samples. Following previously established recommendations, we employed: 1) intensity-based filtering of microarray probes, 2) selection of a representative probe for each gene across both hemispheres, 3) within-donor normalization using the robust sigmoid function and rescaled to the unit interval with the Min-Max function to mitigate donor-specific effects, 4) cross-sample normalization using the same procedure (Arnatkevic iūtė et al., 2019; Markello et al., 2021). This resulted in a final set of six donor-level gene by sample matrices.

### Sparse-to-dense mapping of hippocampal gene expression

*i) Isolating hippocampal samples.* We selected tissue samples falling within the donor-specific *HippUnfold*-derived hippocampal masks and within a 2.5mm radius of the midthickness surface, from both left (*n* = 120) and right (*n* = 58) hemispheres, resulting in 178 samples in total (**Figure 1A** and **Table 1**).
*ii) Surface projection*. Hippocampal samples were mapped to their nearest corresponding vertex on donor-specific surfaces. Given the asymmetrical distribution of samples between hemispheres, samples were mirrored across hemispheres to maximize the number of data points. Expression values of samples assigned to identical vertices were averaged.
*iii) Interpolation of expression values*. We generated spatially dense representations of hippocampal gene expression based on well-known geostatistical method that models spatially varying trends in the data while accounting for spatial autocorrelation among samples: ordinary kriging (Wackernagel, 1995). Briefly, the expression value *Z(s)* at the unsampled location *s* is estimated based on the data from the surrounding locations:

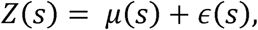

where *μ(s)* is a deterministic trend modeled as a linear combination of spatial coordinates (x, y, z) and □*(s)* is a spatially correlated random field with zero mean and covariance structure *C(h)* defined by an exponential variogram model. For each gene, variogram parameters were estimated empirically from the observed sample distribution.

### Gene expression along the anterior-posterior and medial-lateral axis of the hippocampus

*i) Data-driven transcriptomic associations.* We used partial least squares (PLS) to examine the relationships between gene expression and hippocampal subregional organization, in particular geodesic anterior-posterior and medial-lateral axes of the hippocampus. This multivariate associative technique identifies latent variables maximizing the covariance between two datasets (McIntosh & Lobaugh, 2004). Gene expression profiles and hippocampal maps are correlated with each other, and the resulting covariance matrix is decomposed using singular value decomposition to identify latent variables capturing maximal shared covariance.

The significance of the model was evaluated using non-parametric methods: (*i*) permutation tests were used to assess statistical significance of each latent variable (1,000 permutations; described in **Spatial comparisons** section) and (*ii*) bootstrap ratios were used to assess the stability of gene weights (analogous to *z*-scores, such that 95% and 99% confidence intervals correspond to a bootstrap ratio of ±1.96 and ±2.58, respectively)

*ii) Enrichment analysis*. A web-based gene set analysis toolkit (https://www.webgestalt.org) was used to characterize the biological processes and pathways associated with the significant PLS components (Elizarraras et al., 2024). For a given significant latent variable, genes were ranked according to the magnitude of their loadings and those below a predefined bootstrap ratio threshold (±1.96 for 95% confidence) were excluded from the gene set.

Over-representation analysis determined whether specific processes and pathways were significantly enriched within the selected gene sets. The enrichment ratio was defined as the proportion of overlapping genes between the gene set and ontological category divided by the proportion expected by chance. Statistical significance was assessed using a hypergeometric test.

### Transcriptional hierarchies and its relation to hippocampal multiscale architecture

*i) Organization of gene expression.* We constructed affinity matrices using a normalized angle kernel to measure the similarity of transcriptomic patterns between regions. Diffusion map embedding, a non-linear dimensionality reduction technique implemented in *BrainSpace version 0.1.3* (https://brainspace.readthedocs.io/), was applied to identify low-dimensional eigenvectors that accounted for most of the inter-regional variance. This algorithm uses the diffusion operator *P*α to create a new representation of data and is guided by two key parameters: α, which controls the influence of sampling point density on the manifold, and *t*, which represents the scale of the data. To maintain global connections among data points in the embedded space, α was set to 0.5 (α *=* 0, maximal influence, α *=* 1 no influence), and *t* was set at 0, consistent with previous studies.
*ii) Multiscale contextualization*. Histological (Bieloschowsky silver stain, blockface intensity, calbindin intensity, calretinin intensity, Merker stain, parvalbumin intensity) and MRI-derived (function: first and second principal axes of resting-state functional connectivity; microstructure: fractional anisotropy, mean diffusivity, quantitative T1 intensity, T2* intensity; morphology: curvature, gyrification, thickness) features of the hippocampus were obtained from *HippoMaps version 0.1.0* (https://hippomaps.readthedocs.io/; DeKraker et al., 2025), a toolbox and repository for contextualization of surface-based hippocampal data (**Figure 3B**). To relate the transcriptional organization to multiscale hierarchies of the hippocampus, we computed the correlation between the first and second transcriptomic eigenvectors and each *HippoMaps*-derived map. Permutation test assessed the statistical significance of all associations (1,000 permutations; described in **Spatial comparisons** section)

### Gene contextualization of disease-related hippocampal vulnerabilities

*i) Hippocampal alterations in temporal lobe epilepsy*. We examined transcriptomic associations with well-established markers of hippocampal pathology, namely in pharmaco-resistant temporal lobe epilepsy (TLE). Using multimodal MRI data from 67 patients (mean ± SD age = 31.9 ± 12.0 years; 34 females; 32 left-sided focus) and 92 age- and sex-matched healthy controls (mean ± SD age = 31.7 ± 10.0 years; 51 females), we sampled measures of hippocampal columnar volume (CV) and mean diffusivity (MD; dataset and processing descriptions are provided in the **Supplementary Materials**). Univariate and multivariate alteration maps were defined as vertex-wise *t*-statistics from surface-based linear models comparing patients with TLE to healthy controls while adjusting for age and sex.
*ii) Risk gene associations.* Focusing on *a priori* set of genes associated with TLE (C3orf33, GJA1, KCNAB1, SCN1A, SLC33A1) identified from previous genome-wide association studies (GWAS; Abou-Khalil et al., 2018; Kasperavičiūtė et al., 2013), we correlated each risk gene’s expression with TLE-related alterations. This quantified the extent to which the spatial distribution of each gene aligned with patterns of macrostructural and microstructural changes in patients. Statistical significance was assessed using spatially constrained permutation models (1,000 permutations; described in **Spatial comparisons** section).
*iii) Cell type associations*. Virtual histology is an approach that correlates MRI-derived profiles, such as an interregional profile of group differences in columnar volume and mean diffusivity, with interregional profiles of cell-specific gene expression. Single-nuclei RNA sequencing data from the adult human hippocampus were used to categorize genes as specific to 7 cell types (**Supplementary Table 2**; astrocyte, endothelial, inhibitory neuron, microglia, oligodendrocyte, oligodendrocyte precursor, and pyramidal cells; Ayhan et al., 2021). Cell-specific gene expression was then correlated with hippocampal alterations in TLE to generate a distribution of correlation coefficients for each of the cell types. We assessed the statistical significance of the distribution was assessed using spatially constrained permutation models (1,000 permutations; described in **Spatial comparisons** section).

### Validation of hippocampal gene expression

To assess the robustness of the sparse-to-dense interpolation, we performed a leave-one-out and leave-k-out cross-validation analysis (*k* = 10% to 90% of samples withheld, in 10% increments, 50 iterations). For each iteration, tissue samples were held out, and spatially dense expression maps were reconstructed for all genes using the remaining samples.

### Spatial comparisons

Statistical significance of spatial correlations was assessed using 1,000 eigenstrapping-generated spatial autocorrelation preserving null models, implemented in *HippoMaps* (DeKraker et al., 2025). This framework performs a spectral decomposition of a quantitative map expressed on the hippocampal surface using geometric and then randomly rotate these modes, producing surrogate maps (Koussis et al., 2025).

## Data and code availability

HippoGenes is available in Python and complemented with online documentation (https://hippogenes.readthedocs.io/). Derivative data (https://osf.io/rh2kj/) and code (http://github.io/MICA-MNI/hippogenes) are openly accessible.

## Acknowledgements

A.N. was supported by the Canadian Institute of Health Research (CIHR) and the Fonds de Recherche du Quebec – Santé (FRQ-S). S.L. was funded by CIHR, Centre de Recherche du Centre Hospitalier Universitaire de Sherbrooke (CRCHUS), Université de Sherbrooke, Natural Sciences and Engineering Research Council of Canada (NSERC-Discover, RGPIN-2025-06138), the Fonds de Recherche du Québec – Nature et Technologie (FRQ-NT, 378036) and the Tier-2 Canada Research Chairs Program (CRC). J.R. acknowledged support from NSERC. R.R.C received funding from FRQ-S. M.S. was funded by CIHR. Z.G.O. was supported by the Michael J Fox Foundation, Canadian Consortium on Neurodegeneration in Aging, Baycrest Centre for Geriatric Care, Neuro Genomics Partnership, National Institute of Health (NIH), Silverstein Foundation, Hilary and Galen Weston Foundation and Van Berkhom Foundation. A.B and N.B received funding from FRQ-S, CIHR and Epilepsy Canada. S.A. was funded by NSERC. A.J.B. is a FRQ-S Scholar supported by grants from NSERC. J.W.V. acknowledges funding from the SciLifeLab and Wallenberg Data Driven Life Science Prgram (KAW 2020.0239). S.L.V. received support from the European Research Council Starting Grant Social Connections (SOCO), Hector Research Career Development Award, and Jacobs Canadian Institute for Advance Research (CIFAR) Research Fellowship. M.K. was supported by the Swiss National Science Foundation (219240). R.N.S. is an FRQ-S scholar supported by grants from CIHR, NSERC, and the Alzheimer’s Association. J.D. was funded by NSERC and the Helmholtz International BigBrain Analytics and Learning Laboratory (HIBALL) Association’s Initiative and Networking. B.C.B. acknowledged support from CIHR (FDN-154298, PJT-174995, PJT-206196, CIHR PJT-203761, CIHR PJT-191853), SickKids Foundation (NI17-039), NSERC (RGPIN-2025-05932), Azrieli Center for Autism Research of the Montreal Neurological Institute (ACAR), BrainCanada, FRQS, HIBALL, Healthy Brains and Healthy Lives (HBHL), Centre for Aging + Brain Health Innovation (CABHI), CRC, and the Centre of Excellence in Epilepsy at the Neuro (CEEN).

## Conflicts of interest

J.D.K. and B.C.B. are co-founders of BrainScores and hold stock.

