## Supplementary for "HippoGenes: A robust workflow for subregional transcriptomic profiling of the human hippocampus"

**Supplementary information**

### **Supplementary materials and methods**

#### **Epilepsy case-control dataset**

##### **Participants**

Disease association analyses were conducted on 67 individuals diagnosed with pharmaco-resistant temporal lobe epilepsy (TLE) and radiological evidence of hippocampal sclerosis (HS) and 92 age- (*t* = 0.08, *p* = 0.94) and sex-matched (*χ*² = 0.34, *p* = 0.56) healthy controls. Case-control participants were selected from two independent sites: Universidad Nacional Autonoma de Mexico (EpiC; *n*_TLE/HC_ = 15/25; Rodríguez-Cruces et al., 2020), (*ii*) Montreal Neurological Institute and Hospital (MICs; *n*_TLE/HC_ = 15/36; Royer et al., 2022), and (*iii*) Jinling Hospital (NKG; *n*_TLE/HC_ = 37/31; Weng et al., 2020). Patients were diagnosed according to the classification of the International League Against Epilepsy based on a comprehensive evaluation including clinical history, seizure semiology, video-electroencephalography recordings, neuroimaging, and/or neuropsychological assessment. We excluded patients who had encephalitis, a history of traumatic brain injury, or bilateral TLE diagnosis. At the time of data analysis, all patients had standard radiologically suspected hippocampal sclerosis (HS) from preoperative examination, with 46 having undergone resective surgery (40 had undergone cortico-amygdalo-hippocampectomy and 6 had undergone selective amygdalo-hippocampectomy). At post-surgical follow-up (mean ± standard deviation [SD] = 2.48 ± 2.18 years), 34 patients have been seizure-free (Engel I), 5 have shown significant reductions in seizure frequency (Engel II), 4 have shown prolonged seizure intervals (Engel III), 2 have shown no worthwhile improvements (Engel IV), and 1 was lost for follow-ups. Based on established histopathological criteria, all available specimens showed HS or gliosis. Data collection was approved by the ethics committees of each site, and all participants provided written informed consent in accordance with the Declaration of Helsinki.

##### **MRI acquisition**

Every participant underwent a research-dedicated multimodal MRI scan before surgery. Data from EpiC were collected on a 3T Philips Achieva TX scanner equipped with a 32-channel head coil and included: (*i*) high-resolution T1-weighted (T1w) MRI using a 3D spoiled gradient-echco (voxel size = 1×1×1mm^3^, repetition time [TR] = 8.1ms, echo time [TE] = 3.7ms, flip angle [FA] = 8°, inversion time [TI] = 900ms, field of view [FOV] = 256×256mm^2^) and (*ii*) diffusion MRI imaging using 2D echo-planar imaging with 2000s/mm^2^ and 60 diffusion directions (voxel size = 2×2×2mm^3^, TR = 11860ms, TE = 64.3ms, FOV = 256×256mm^2^).

Data from MICs were collected on a 3T Siemens Magnetom Prisma-Fit scanner equipped with a 64-channel head coil and included: (*i*) high-resolution T1w MRI using a 3D magnetization-prepared rapid gradient-echo sequence (MPRAGE; voxel size = 0.8×0.8×0.8mm^3^, matrix size = 320×320, repetition time [TR] = 2300ms, echo time [TE] = 3.14ms, flip angle [FA] = 9°, inversion time [TI] = 900ms, field of view [FOV] = 256×256 mm^2^), (*ii*) diffusion MRI using a 2D spin-echo echo-planar imaging sequence with multi-band acceleration consisting of three distinct shells with b-values 300, 700, and 2000 s/mm^2^ and 10, 40, 90 diffusion weighting directions, respectively (voxel size = 1.6×1.6×1.6mm^3^, TR = 3500ms, TE = 64.40ms, FA = 90°, FOV = 224×224mm^2^).

Data from NKG were collected on a 3T Siemens Trio scanner with a 32-channel head coil, and included: (*i*) high-resolution T1w MRI using a 3D-MPRAGE sequence (voxel size = 0.5×0.5×1mm^3^, TR = 2300ms, TE = 2.98ms, FA = 9°, FOV = 256×256 mm^2^, 176 slices) and (*ii*) diffusion MRI using a spin echo-based echo planar imaging sequence consisting of four distinct shells with b-values of 1000 s/mm^2^ (voxel size = 0.94×0.94×3mm^3^, TR = 6100ms, TE = 93ms, FA = 90°, FOV = 240×240mm^2^).

##### **MRI processing**

Multimodal MRI data were processed using micapipe version 0.2.3 ( <https://micapipe.readthedocs.io/>; Cruces et al., 2022) and *HippUnfold version 1.3.0* (<https://hippunfold.khanlab.ca/>; DeKraker et al., 2022).

T1w MRI data were deobliqued, reoriented to standard neuroscience orientation (LPI: left to right, posterior to anterior, and inferior to superior), corrected for intensity nonuniformity, intensity normalized, skull-stripped, and used to extract surface models of the hippocampus. Subject-specific columnar volume was measured as the multiplication of hippocampal thickness (Euclidean distance between corresponding inner and outer layer vertices) and surface area (one third of the area of each triangle a vertex is a part of).

Diffusion MRI data were preprocessed using MRtrix3 and included denoising, b0 intensity normalization as well as correction for susceptibility distortion, head motion, and eddy currents. Diffusion tensor-derived mean diffusivity (MD), representative of tissue microstructure), was sampled and interpolated along the hippocampal surface.

##### **Multisite data harmonization**

##### Morphological and microstructural data were harmonized across sites using Combat (<https://github.com/Jfortin1/ComBatHarmonization>), a post-acquisition statistical batch normalization to harmonize between-site effects while preserving effects of age, sex and disease status (Fortin et al., 2018)

### **Supplementary figures**

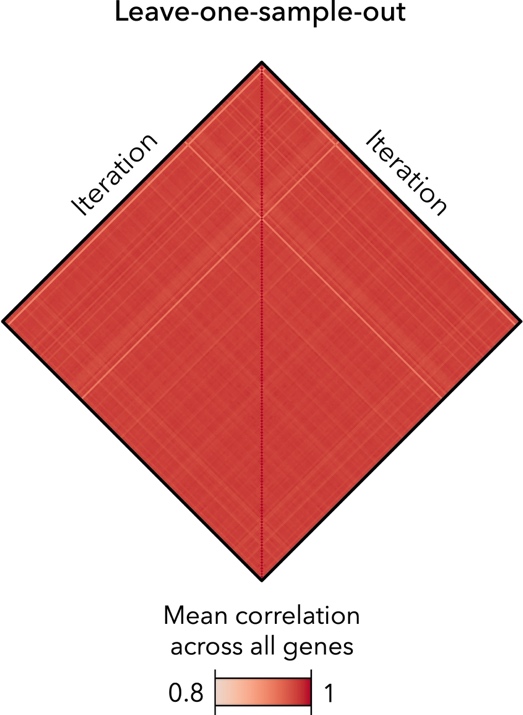

**Supplementary figure 1 |** Leave-one-sample-out cross validation of hippocampal gene expression profile showing high similarity between iterations.

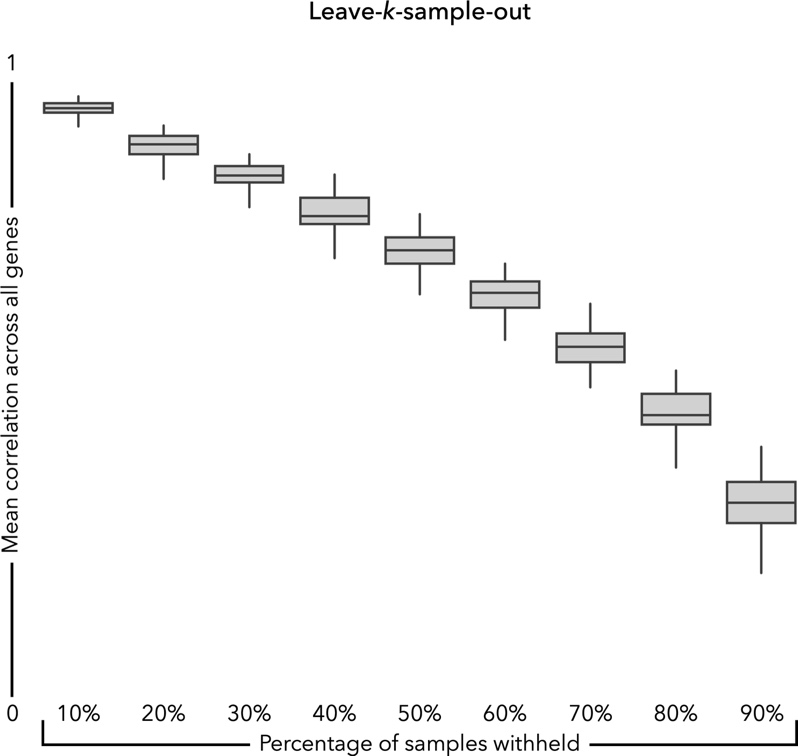

**Supplementary figure 2 |** Leave-*k*-sample-out cross validation of hippocampal gene expression profiles. Boxplots represent the mean correlation across all genes between empirical (see *Figure 1*) and reconstructed atlases (from 10% to 90% samples withheld, 10% increments), ends of boxplots represent the first (25%) and third (75%) quartiles, the centre line (median) represents the second quartile of the 50 cross-validation iterations, the whiskers represent the non-outlier endpoints of the distribution

### **Supplementary tables**

**Supplementary table 1 |** Summary statistics (mean, standard deviation and range of correlations) of leave-*k*-sample-out cross validation of hippocampal gene expression profiles.

| Samples withheld | Mean | Standard deviation | Range |
| --- | --- | --- | --- |
| 10% | 0.915 | 0.015 | 0.874 – 0.936 |
| 20% | 0.862 | 0.018 | 0.812 – 0.892 |
| 30% | 0.816 | 0.020 | 0.768 – 0.850 |
| 40% | 0.762 | 0.028 | 0.694 – 0.819 |
| 50% | 0.705 | 0.029 | 0.639 – 0.760 |
| 60% | 0.637 | 0.030 | 0.554 – 0.685 |
| 70% | 0.560 | 0.033 | 0.458 – 0.626 |
| 80% | 0.462 | 0.034 | 0.379 – 0.525 |
| 90% | 0.324 | 0.044 | 0.222 – 0.411 |

**Supplementary table 2 |** Cell-type specific gene profiles from Ayhan et al., 2021.

| Cell | Genes |
| --- | --- |
| Astrocyte | ABLIM1, ACACB, ACOT11, ACSS1, ACSS3, ADAM10, ADAMTS19, ADAMTS9, ADCY2, ADD2, ADGRA3, ADGRV1, ADSS, AHCYL1, ALDH1A1, ALDH1L1, ANKRD36, ANKRD44, APBA1, AQP4, ARHGEF4, ARMC8, ASPH, ATE1, ATP13A4, ATP1A2, ATP1B3, BACH1, BBS9, BBX, BCL2L1, BMPR1B, BTBD7, C1orf61, C21orf91, C9orf3, C9orf85, CABLES1, CACNA1B, CACNA1C, CALN1, CAMK2A, CAMK2G, CCDC146, CCDC91, CD44, CDC42BPA, CDH20, CHPT1, CLIP4, CLU, COL5A3, CPEB1, CPNE2, CPNE4, CPPED1, CRYAB, CTDSPL, CTNNA2, CTNND2, CUEDC1, CYFIP1, CYLD, DAPK2, DCLK2, DENND5A, DIP2C, DPP10, DTNA, DTNB, EEF2K, EFHD1, ELF1, ELOVL7, EML2, EPHA6, EPHA7, ERBB3, ETNPPL, EYA1, FAM126A, FAM13B, FAM153C, FAM184A, FAM189A2, FAM196B, FARP1, FAXDC2, FBXW7, FGFR3, FKBP5, FLNB, FMN2, FTCDNL1, GAB2, GABRA1, GADD45G, GAN, GATS, GFAP, GLCCI1, GLCE, GLI3, GLIS3, GPC5, GPM6A, GPR75-ASB3, GRAMD3, GRB10, GRIN1, GRIP1, GTDC1, HAPLN2, HID1, HIF3A, HPSE2, HSP90AA1, HTR4, IFLTD1, INSR, IQCA1, ITCH, ITPKB, ITPR1, KAT2B, KCNB2, KCND2, KCNH7, KCNN3, KIF5C, KLHL3, KLHL42, LCOR, LHFP, LRIG1, LRRC1, LRRC16A, LRRFIP2, LY6H, MAN1A1, MAP3K5, MAPK4, MARCH3, MBNL1, MFSD6, MGAT4C, MICAL3, MIPEP, MLH3, MRVI1, MSI2, MT2A, MTMR7, NALCN, NBEA, NDFIP2, NEBL, NEGR1, NEK3, NETO2, NFATC2, NIPAL3, NKAIN3, NPL, NR2C2, NSMCE2, NTNG1, NTRK2, NYAP2, OLFM3, OSBP2, PACS2, PADI2, PAMR1, PARD3, PARD3B, PCBP3, PDE7B, PGBD5, PHC2, PIK3C2B, PITPNC1, PLA2G4C, PPP2R2A, PPP2R2B, PPP2R3A, PPP4R2, PRDM2, PREX1, PREX2, PRKACB, PRKCG, PRKG1, PRODH, PTK2B, PTP4A2, RAB11FIP4, RABEP1, RANBP3L, RAP1GDS1, RBFOX3, RFTN2, RFX4, RHOBTB1, RHOU, RIMBP2, RNF144A, RORA, RP11-281A20.3, RP11-566D24.2, RUNDC3B, RYR3, SAMD3, SCFD2, SCN3B, SDK1, SHROOM3, SIPA1L2, SLC14A1, SLC1A2, SLC22A23, SLC25A18, SLC35F1, SLC38A9, SLC4A4, SLC4A7, SNCA, SNX1, SNX29, SNX30, SNX9, SORBS1, SOS1, SOX5, SPARCL1, SPATA6, SPECC1, SRRM4, STK33, STON2, STXBP3, TBC1D4, TET3, TFEB, TFRC, TLE1, TMEM117, TMEM181, TMEM2, TMEM64, TMTC4, TNC, TNIK, TNPO1, TPD52, TPD52L1, TRPM3, TRPS1, TSPAN5, TUBA1B, TUBB4A, UBE2R2, ULK2, UNC13C, URI1, USP15, USP31, WASF1, WDR49, WNK1, XYLT1, ZCCHC6, ZFP36L1, ZNF148, ZNF331, ZNF385B, ZNF608, ZNRF3 |
| Endothelial cell | ABCB1, ABLIM1, AFF1, AKAP12, AP2M1, AP3S1, ARPP19, ATP10A, ATP1A1, ATP2A2, ATP2B1, CALM3, CAP1, CCNL1, CD55, CLDN5, CNOT6L, CNR1, COL5A2, CSRP1, CTNNA1, CYTH3, EFNB2, EGR3, EIF4A2, ELOVL5, EML4, EPB41L4A, ERC1, FERMT2, FLNB, FLT1, GABARAPL1, GALNTL6, GBE1, GUK1, HINT1, HNRNPA2B1, HNRNPK, HNRNPLL, HNRNPU, HSPA1A, IFNGR1, ITPR1, KANSL1L, LAMP1, LDLRAD3, LONRF1, LTBP1, MAPK1IP1L, MARCH3, MECOM, MYO1E, NAP1L1, NHSL1, NR4A2, NUP58, OPCML, PABPC1, PDE5A, PELI1, PELI2, PIAS1, PID1, PIP5K1B, PITPNB, PRKAR1A, PRNP, RAPGEF1, RGS6, RHEB, RNASE1, RNF19A, RUSC2, SARAF, SEC14L1, SERINC1, SIDT1, SLC4A7, SNAP25, SOD1, SQSTM1, SRGAP2, SRSF3, SRSF7, STK38, TACC1, TBC1D1, TMOD3, TOP1, TRA2B, TTC28, TUBA1A, VPS37B, VSNL1, WLS, YWHAB, YWHAG, ZIC1, ZNF385D |
| Inhibitory neuron | ABHD6, ACSL1, ACTG1, ACYP2, ADAM10, ADAMTS17, ADD3, ADSS, AFMID, AGO2, AHCYL2, ANKIB1, ANKRD10, ANKRD28, ANKRD44, ANXA4, APLP1, APOD, ARGLU1, ARHGAP23, ARHGEF2, ARID2, ATG10, ATG13, ATL1, ATP11A, ATP1B3, ATP2B1, ATP2B4, ATP8B4, B2M, BCAS3, BIN1, BPTF, C12orf76, C14orf145, C16orf88, C8orf46, C9orf85, CADPS2, CALD1, CCBE1, CCDC146, CCNY, CDK5RAP2, CDYL, CGGBP1, CLU, CNOT2, COL16A1, COL5A2, CPEB2, CPEB4, CPPED1, CRADD, CREB3L2, CREG2, CST3, CTBP2, CTNNA1, CTSB, CTTNBP2, CUEDC1, CYFIP1, CYP7B1, DECR1, DOCK9, DPYSL2, DTWD1, EFCAB11, EGR1, EIF3E, ELL2, EMC2, EML1, ENC1, ENDOD1, ETNK1, EXOC4, EXOC6, FAM102A, FAM134B, FAM13A, FAM149A, FANCL, FAR1, FBXO32, FBXW7, FCHSD2, FEZ1, FGF1, FGFR1OP2, FOXP1, FTCDNL1, GAB2, GCLC, GPBP1, GPCPD1, GPM6A, GREB1L, HBP1, HBS1L, HIBCH, HIF1A, HNRNPA2B1, HNRNPDL, IVNS1ABP, IWS1, KCNK1, KCNT2, KCTD8, KMT2C, KRCC1, LANCL1, LDB2, LPP, LRRN1, LYPLAL1, MAP3K5, MAP4, MARCKSL1, MAST3, MAT2A, MED13L, MGAT5, MLLT4, MMD2, MOB1B, MON2, MT2A, MTERF1, MTURN, MXI1, MYO5A, MYO9B, NDFIP2, NDRG2, NFE2L2, NIPAL3, NR3C2, NRIP1, NSRP1, NTNG1, NUMB, NXPE3, OLIG1, OXR1, PAG1, PAK2, PAPD4, PARD3, PAXBP1, PCBP3, PDCD6IP, PDGFC, PHF21A, PHIP, PIGF, PIK3C2B, PIK3R1, PKP4, PLEKHA1, PLEKHA2, PLEKHG1, POGZ, POLK, PPFIA2, PPM1H, PPP1R12A, PPP1R9A, PPP4R2, PPP4R3B, PRTFDC1, PSME4, PTP4A2, PTPN11, PTPRF, RAB10, RASGRF2, RASSF8, RBM26, REEP3, REL, RGS5, RHBDL3, RHOBTB1, RHOBTB3, RILPL1, RIT2, RNASE1, RNF130, RNF213, RP1-5O6.7, RP11-2E17.1, RP11-437B10.1, RP11-717K11.2, S100PBP, SAMD3, SCARB2, SEPT4, SERPINI1, SF1, SIDT1, SLC11A2, SLC35F3, SLC38A9, SLC48A1, SLC4A4, SLF2, SLTM, SMURF2, SNX1, SORCS2, SOX2, SPAG9, SPG20, SPRED1, SRRM2, SS18, STK4, STPG2, SUMF1, TBL1XR1, TCP11L2, THOC3, TLE1, TMED10, TMEM59, TMEM87A, TPCN1, TRA2A, TRPC4AP, TSC22D2, TSC22D4, TTC7A, TUBA1A, TUBB4A, UBB, UBE2R2, UNC5C, USP3, USP39, USP40, USP47, UST, UVRAG, VAPA, VMP1, WAPL, WASHC5, WDFY2, WDR20, WSB1, XRCC4, YPEL2, ZBED5, ZBTB44, ZC3H6, ZFAND6, ZKSCAN1, ZMIZ1, ZNF146, ZNF397, ZNF521, ZNF644, ZZZ3 |
| Microglia | ABCC4, ABHD12, ABR, ACAP2, ADAM28, ADRBK2, AFTPH, AKAP13, ANAPC16, ANKRD44, AOAH, APBB1IP, ARHGAP15, ARHGAP22, ARHGAP24, ATM, ATP2A2, ATP6V0A1, ATP8B4, B3GNT5, B4GALT1, BASP1, BMP2K, BNC2, BRD4, C10orf11, C15orf41, C3, CAMK2D, CAP1, CCL3, CCND3, CD74, CD83, CD86, CDK6, CHKA, CHST11, CREBBP, CRK, CROCC, CSGALNACT1, CTNNB1, CYFIP1, DENND4A, DISP1, DNM2, DOCK2, DOCK8, DSCAM, E2F3, EP300, EPS8, ERGIC1, FAM117B, FAM149A, FAR2, FBXW11, FGD4, FKBP5, FLI1, FMN1, FOS, FOXN3, FOXP2, FRMD4A, FUS, FYB1, GAK, GALNT1, GAPVD1, GNG7, GPD2, GRAMD1B, HEATR5B, HERPUD1, HIF1A, HNRNPD, HS3ST4, HSP90AA1, IGSF21, IKZF1, IL1RAP, INPP5D, IPCEF1, IRAK2, IRAK3, ITPR2, ITSN1, KDM2A, KDM3B, KDM4B, LDLRAD4, LHFPL2, LIMS1, LPCAT2, LRRK1, LYN, MAML3, MAN1A1, MB21D2, MCF2L2, MEF2A, MEF2C, MERTK, MGMT, MPRIP, MRC1, MTDH, MYO1E, MYO1F, NFIC, NFX1, NHSL1, OLR1, P2RY12, P4HA1, PACS1, PACSIN2, PAG1, PALD1, PAN3, PCNX2, PDE3B, PDGFC, PELI2, PER1, PIK3R1, PIK3R5, PIP5K1B, PITPNA, PITPNB, PLA2G4A, PLCL2, PLXDC2, PPP6R2, PRKCH, PRMT2, PRRC2B, PSMA1, PTK2B, PTPRC, PTPRE, QSER1, RAB10, RAB31, RAF1, RAP1B, RAP1GAP2, RASGEF1C, RB1, RBFOX1, RBM47, RCOR1, RHBDF2, RHEB, RIN3, RNF145, RNF149, RNF216, RUNX1, SCAF11, SDK1, SEC14L1, SEPT2, SERPINE1, SESN3, SETD3, SFMBT2, SH3TC1, SLC11A1, SLC1A3, SLCO2B1, SMAP2, SNCA, SNX29, SNX9, SOCS6, SPP1, SPTLC2, SRGAP2, SRGAP2B, SRGAP2C, SRGN, SRSF4, ST6GAL1, ST6GALNAC3, STAG1, STAU1, STIM1, SUSD6, SYK, SYNDIG1, TAB2, TANC1, TBC1D16, TBXAS1, TET2, TET3, TGFBR2, THADA, TLR2, TM9SF3, TMCC3, TMEM156, TNFRSF1B, TPM3, TRAPPC9, UBASH3B, UBL3, USP24, USP25, USP53, UVRAG, VOPP1, VPS13D, WDR37, WDR70, XPR1, ZFAND3, ZFHX3, ZNF106 |
| Oligodendrocyte | AATK, ABCA2, ABCA6, ABCA8, ABTB2, AC109326.1, ACAP3, ACTB, ADAMTS18, ADCY3, ADD1, ADGRE2, ADIPOR2, AFMID, AGPAT4, AMER2, ANK3, ANLN, ANO4, APLP2, ARAP2, ARFGAP1, ARHGAP1, ARHGAP23, ARNT2, ATP10B, BAIAP2L2, BCAS1, BCL6, BIN1, BTBD17, C10orf90, C1QTNF3-AMACR, CALD1, CARNS1, CBR1, CBWD5, CCDC88A, CCP110, CD22, CDK18, CDK19, CERCAM, CHRM5, CIRBP, CKB, CLIP2, CLMN, CNDP1, CNP, CNTN2, CNTNAP4, COL16A1, COL18A1, COL24A1, CPOX, CREB5, CREM, CSRP1, CTNNA3, CUEDC1, DAAM2, DAPK2, DDX17, DENND2A, DIP2B, DLC1, DLG1, DNAJB2, DNAJC6, DNM3, DOCK5, DPYD, DPYSL5, DSCAML1, EDIL3, EFHD1, EGR1, EHBP1, ENOX1, ENPP2, ENPP6, EPS15L1, ERBIN, EVA1C, EXOC6B, FA2H, FAM107B, FAM124A, FAM13C, FMN1, FMNL2, FOLH1, FOS, FRMD4B, FRMD5, FRYL, FUT8, GAB1, GFAP, GIPC1, GNA12, GNA13, GNG7, GRM3, GSN, HAPLN2, HDAC9, HECTD2, HEPACAM, HHIP, HID1, HIF1A, HIP1, HMGCS1, HNRNPL, HS3ST4, HS3ST5, INF2, IRS2, ITGA2, JUND, KANK1, KANK4, KCNAB1, KCNH8, KCNMB4, KCNN2, KIAA0930, KIAA1755, KIF13B, KIF19, KIF6, KLHL32, LAMA2, LDB3, LDLRAD4, LENG8, LGR5, LINGO1, LMF1, LMO4, LPAR1, LPCAT2, LPGAT1, LPIN1, LRP12, LRP2, LRRC1, LRRC63, MAL, MAN2A1, MAP4K4, MAP7, MARCH1, MAST4, MBP, MDM4, MED13L, MGEA5, MKRN3, MOB3B, MOBP, MOG, MT2A, MT3, MTUS1, MVB12B, MYO1D, MYO9B, MYRF, NAMPT, NASP, NCKAP5, NCOA7, NDE1, NDRG1, NDRG2, NDST3, NDUFA10, NINJ2, NME3, NPC1, NR4A2, NRG1, NUMA1, OLFM3, OPALIN, OTUD7A, PABPN1, PALM2, PCBP4, PCDH9, PCSK6, PDE1A, PDE1C, PDE4B, PDE8A, PEX5L, PHLDB1, PHLPP1, PIEZO2, PIK3C2B, PIP4K2A, PLA2G16, PLCL1, PLD1, PLEKHH1, PLLP, PLXDC2, PLXNB1, POLR2F, PPP1R12A, PPP1R14A, PPP1R16B, PPP2R2B, PRKCB, PRKD1, PRR5L, PRUNE2, PSEN1, PTPRK, PXK, QDPR, QKI, QTRT1, RAPGEF3, RASGRF2, RASSF4, RBM5, RFTN2, RGS12, RIN2, RNF130, RNF220, ROGDI, ROR1, RP11-654K19.6, RP11-792A8.5, SAMD4B, SCD, SEC14L5, SELO, SEMA3B, SEPT7, SF1, SGCD, SGK1, SH3GL3, SH3GLB2, SH3TC2, SHTN1, SIK3, SLAIN1, SLC22A15, SLC24A2, SLC27A1, SLC44A1, SLC5A11, SLCO1A2, SORCS2, SPOCK1, SPOCK3, SRCIN1, SRRM2, ST18, STAT2, STMN4, SUN2, SVEP1, SYNJ2, TCF7L2, TECPR2, TF, TFEB, TMC6, TMC7, TMEFF2, TMEM144, TMEM165, TMEM235, TMEM259, TMEM63A, TMSB10, TMTC2, TP53TG5, TPPP, TPST1, TRAK1, TRIM2, TTC3, TTLL7, TTYH2, UGT8, UNC5C, USP39, VRK2, XPR1, ZDHHC20, ZFYVE16, ZNF385D, ZNF536, ZNF565, ZNF652 |
| Oligodendrocyte precursor cells | ABAT, ABCG1, ABHD2, AC116366.7, ACSL3, ACSS2, ADAMTS17, ADCY3, ADCY5, ADCY8, ADSS, AFAP1L2, AHI1, AKAP13, AMZ1, ANAPC5, ANKFN1, ANKRD12, ANKRD36, ANKRD36C, ANTXR1, APBA2, APP, ARHGAP31, ARHGAP44, ARHGAP5, ASCC1, ATCAY, ATP13A4, ATP1B1, ATRNL1, BACH2, BBX, BORCS5, BRINP3, BRMS1L, C11orf49, C16orf62, C1orf198, C21orf91, C6orf62, C9orf85, CA10, CACNA1A, CACNA1D, CACNG4, CAMK2B, CAMSAP2, CBFA2T2, CCDC136, CCDC50, CCDC64, CCND3, CHCHD6, CHL1, CHN1, CHST11, CHST9, CLIP4, CLVS2, CMYA5, CNKSR3, CNTN2, CNTNAP3B, CNTNAP5, COL11A1, COL4A3, COL9A1, CPNE2, CPNE4, CRB1, CREM, CRISPLD2, CRYL1, CSMD1, CSMD2, CTD-2561J22.3, CTNNBIP1, CUEDC1, CUX1, CYFIP2, CYP27A1, DCC, DGKG, DGKZ, DHRS7B, DIP2C, DIS3L2, DISC1, DKK3, DNAH7, DNAJB2, DNASE1, DOCK7, DOK5, DPP6, DSCAM, DTNB, EEF2K, EFR3B, EGFR, EPHA5, EPN2, ERGIC1, ETV5, EVL, EYA3, FAM110B, FAM126A, FAM134B, FAM149A, FAM153C, FAM171B, FAM196B, FAM78B, FAR1, FARP2, FARS2, FBN1, FBXL2, FEZ2, FGF12, FGF14, FGFR1, FNIP2, FOXN3, FSD1L, GABRG2, GALNT13, GAN, GATM, GFRA1, GNB1, GOLGA8A, GPC6, GRAMD1B, GRB10, GRID2, GRIK1, GRIN1, GSG1L, GTDC1, HACD2, HACL1, HAPLN2, HECW1, HERPUD1, HRASLS, HSPA1A, HYDIN, ITGA8, ITPK1, ITPKB, KANSL1L, KAT2A, KAT2B, KAZN, KCNB2, KCND2, KCNH1, KCNK1, KCNN3, KCNT2, KIAA1370, KIAA1468, KIAA1549L, KIF1B, KIF26B, KIF5C, KLHL3, KSR2, LGI1, LHFPL3, LIMD1, LMNA, LMO7, LRFN5, LRP1, LRRC4C, LRRTM4, LUZP2, MAP1B, MAP1LC3A, MAP3K14, MAP3K3, MAP3K5, MAPT, MBOAT2, MDGA2, ME1, MEGF11, MEGF9, MFAP3L, MGLL, MKNK1, MLH3, MMD2, MMP16, MPC1, MPHOSPH8, MPPED2, MTSS1L, MTURN, MVB12B, MYO10, MYO1E, MYT1, N4BP2L1, NAALAD2, NDFIP2, NDRG3, NIM1K, NIPAL3, NLK, NOP56, NOVA1, NPL, NR1D2, NRBP2, NRCAM, NRXN2, NSMCE2, NTNG1, NUP98, NXPH1, NYAP2, OPCML, OSBPL6, PAAF1, PACSIN2, PALLD, PARP8, PBX1, PCDH15, PCDH7, PCM1, PCSK5, PDS5B, PDZD2, PDZRN4, PELI2, PER1, PHACTR4, PHC3, PHLDB1, PID1, PIGN, PIK3CB, PKD1, PLEKHA2, PLEKHG1, PLEKHM3, PLPP1, PLPP4, PMS1, POGZ, POU6F2, PPFIBP2, PPP3CA, PRKACB, PRKCB, PRKCG, PRKCZ, PTBP2, PTPN13, PTPRG, PTPRT, PTPRZ1, RAB10, RAB11FIP4, RABEP1, RABGAP1, RAF1, RALGPS2, RANBP17, RAP1GDS1, RBM26, RBMS1, RBMS3, RCOR1, RGL1, RGS12, RHBDL3, RMND5A, RNF157, RNF180, RNF19A, RNF24, ROGDI, RORA, RPRD2, SAMD4B, SCMH1, SCN1A, SCN8A, SEPP1, SEPT4, SETD7, SEZ6L, SGCZ, SH3PXD2B, SIK2, SLC11A2, SLC1A2, SLC2A13, SLC35F1, SLC8A3, SMAD3, SMOC1, SNAP91, SNED1, SNTG1, SORCS1, SORCS3, SOX5, SOX6, SREBF2, STARD9, STAT3, STK32A, STK33, STOX2, SULF2, TANC1, TCP11L2, TFEB, THSD7A, TIMM23B, TLN2, TMEM132C, TMEM132D, TMEM232, TNK2, TNR, TNS3, TOB2, TPM1, TRAK1, TRERF1, TSEN15, TTLL5, TUBB4A, USP32, USP47, UST, VCAN, WDR59, XKR4, XYLT1, YAF2, YWHAQ, ZC3H13, ZDHHC14, ZEB1, ZFYVE16, ZFYVE28, ZNF189, ZNF365, ZNF462, ZNF483, ZNF532, ZNF565, ZNF609, ZNF714, ZNF75A, ZNF827, ZRANB3, ZSWIM5, ZZZ3 |
| Pyramidal neuron | ABCG1, ACSS2, AGO3, AGPAT4, AGPS, AKAP10, ANKS1A, ANXA4, ARHGAP5, ATAD2B, ATE1, ATF7IP, BCL2L1, BTBD3, BTBD8, C16orf88, C3orf58, CAMK2N1, CAST, CCM2, CD81, CDK13, CDK17, CDK5RAP2, CDYL, CEP170, CFTR, CHPT1, CHSY1, CLU, COL18A1, COL4A3BP, CPOX, CREM, CTDSPL2, CTNNA1, CUEDC1, CYTH1, DDX5, DECR1, DMXL1, DPP8, DPY19L3, EGR3, EIF4E, EMC2, ENDOD1, EPB41L4A, ETNK1, FAM110B, FAM169A, FAM196B, FAR1, FBXL4, FBXO7, FGF1, FNDC3A, FTH1, GALNT10, GALNT7, GCFC2, GNA14, GOLGA4, GOLPH3, GPBP1, HACD2, HERPUD1, HIBADH, HMGCS1, HMGN3, HNRNPC, HSD17B12, HSD17B4, HSPA8, INO80D, ITCH, KCNN3, KIAA1109, KIDINS220, KLF13, KLF9, KLHL20, KRIT1, LDB3, LDLRAD3, LRRTM4, LYPLAL1, MANBA, MAP4, MAP4K3, MAT2A, MFN1, MGAT4A, MGAT5, MLLT4, MPDZ, MSL2, MTDH, MYO18A, NCOA2, NECAB1, NF1, NFASC, NFKB1, NHLRC3, NRIP1, NSD1, NXPE3, OSBPL11, OSBPL3, OSTF1, OTUD7A, PCDH17, PDGFC, PGAP1, PHACTR3, PHC3, PIGU, PKP4, PPP1R9A, PPP2R5A, PRKACB, PSMA1, PTAR1, PTP4A2, PUM1, RAB11FIP2, RBBP4, RCBTB1, RCOR3, RECK, RGS5, RHBDD1, RHEB, RHOA, RHOBTB1, RNF170, RP11-2E17.1, RPL41, RUFY2, S100PBP, SAMD4A, SAMD4B, SEC14L1, SEPT8, SF1, SH3GLB1, SHPRH, SLC12A6, SLC38A2, SLC38A9, SLC4A4, SNAP23, SNX29, SNX30, SNX6, SORT1, SP3, SPOP, SRSF10, STAT3, STOX2, TARSL2, TBL1XR1, TET2, TMED10, TMEM260, TMEM64, TNKS, TOGARAM1, TPD52, TPM3, TPPP, TSEN15, TTC27, TTYH1, TULP4, UBA3, UBE2E1, USP47, WAC, WASHC3, WDR41, WNK1, XRCC4, XRCC5, YPEL2, YTHDC1, ZBED5, ZCCHC6, ZDHHC14, ZHX2, ZMYM2, ZNF207, ZNF331, ZNF654 |
